# Systematic mapping of insertion-tolerant regions enables capsid engineering of the RNA phage PP7

**DOI:** 10.64898/2026.04.15.718622

**Authors:** Shin-Yae Choi, Min-Ju Kim, Oh Hyun Kwon, Chanseop Park, Hee-Won Bae, Hongbaek Cho, You-Hee Cho

**Affiliations:** Program of Biopharmaceutical Science, Department of Pharmacy, College of Pharmacy and Institute of Pharmaceutical Sciences, CHA University, Gyeonggi-do 13488, Korea; Department of Biological Sciences, College of Natural Sciences, Sungkyunkwan University, Republic of Korea

**Author notes:** Shin-Yae Choi, Min-Ju Kim, and Oh Hyun Kwon contributed equally to this work.

**Keywords:** RNA phage, insertion-tolerant regions, synthetic phages, capsid tagging

## Abstract

RNA phages are attractive platforms for the design of programmable bioparticles, but their development has been constrained by limited knowledge of genomic sites that can tolerate sequence insertion. Here, we combined MuA transposase-mediated in vitro insertion mutagenesis with our established reverse genetics systems to systematically identify insertion-tolerant regions (ITRs) in the RNA phages MS2 and PP7. Screening of 4,555 MS2 and 2,228 PP7 random insertion clones identified 29 and 26 nonredundant ITRs, respectively. We further analyzed and compared these ITRs in the context of RNA genome organization and virion architecture. Both phages contained ITRs within the maturation protein (MP), whereas only PP7 tolerated insertions within the coat protein (CP). On the basis of structural location and plaque-forming capacity, an ITR situated between Gly74 and Glu75 (GGC^GAG) in the PP7 CP was selected for further study. Infectious phage particles generated from cDNA clones retained the 15-bp insertion at both the RNA and protein levels. Engineered PP7 phages carrying an RGD motif inserted into the CP at this ITR displayed enhanced in vivo clearance in a *Drosophila* model, despite having in vitro stability comparable to that of the wild type. These findings provide the first example of CP engineering in an RNA phage and establish a framework for engineering RNA phages for biological and biotechnological applications.

**IMPORTANCE:** A major obstacle to developing RNA phages as synthetic biology platforms is the lack of design principles for genomic insertion. Here, we address this limitation by establishing a mutagenesis-and-recovery workflow that systematically identifies insertion-tolerant regions (ITRs) in the RNA phages MS2 and PP7. The resulting maps reveal distinct structural constraints in the two phages and enable rational engineering of a peptide-display site in the PP7 capsid. Using this approach, we generated an engineered infectious phage with a modified capsid, thereby providing the first demonstration of capsid engineering in an RNA phage. This study lays the groundwork for the rational design of RNA phage virions as tractable and engineerable scaffolds for future biological and biotechnological applications.

## INTRODUCTION

Single-stranded RNA (ssRNA) bacteriophages of the family *Fiersviridae*, including *Emesvirus zinderi* (MS2), *Qubevirus durum* (Qβ), and *Pepevirus rubrum* (PP7), are among the simplest yet most informative biological entities. These ssRNA phages exemplify fundamental properties that are also central to eukaryotic RNA viruses, particularly the coupling between RNA structure and function (1, 2) and the rapid evolution driven by high mutation rates (3, 4). In the current era of RNA-based biopharmaceuticals, these viruses serve as paradigmatic model systems for RNA biology and biotechnology (5-7). Their inherent attributes characterized by pronounced capsid uniformity, a sophisticated capacity for selective RNA packaging, and overall structural simplicity, make them attractive candidates for development as synthetic chassis for RNA delivery and molecular display (6, 8, 9).

Despite this promise, progress in RNA phage biotechnology has been limited by a critical bottleneck: the lack of robust genome-wide engineering strategies that preserve the multilayered informational density of the viral genome (10-12). The ssRNA genome is a highly compact evolutionary product in which a single sequence frequently fulfills multiple functions, including protein coding, RNA secondary structure formation, and virion assembly orchestration. Consequently, even minor perturbations often abolish viral viability or compromise capsid integrity. To mitigate these pleiotropic effects, it is therefore imperative to identify and characterize specific genomic positions, termed insertion-tolerant regions (ITRs), that can accommodate heterologous sequence insertions without deleterious consequences, analogous to those identified in certain eukaryotic RNA viruses (13-15). Although MS2 and PP7 coat-protein dimers are well established scaffolds for peptide display and cargo conjugation (8, 16, 17), comprehensive ITR maps for these phages remain lacking. Such maps are an essential prerequisite for programmable genome engineering and would facilitate both fundamental studies of RNA phage biology and practical engineering of RNA phages. In this study, we performed a systematic, genome-wide analysis of ITRs in two representative fiersphages, MS2 and PP7, exploiting an in vitro transposon-mediated mutagenesis strategy. By screening large libraries of transposon-inserted variants, we generated comprehensive ITR profiles and uncovered distinct insertion tolerance landscapes in the two phages. Leveraging a newly identified ITR within the capsid-coding region, we successfully engineered chimeric PP7 particles displaying the Arg-Gly-Asp (RGD) peptide motif (18). We further assessed the functional consequences of this capsid modification by demonstrating altered in vivo clearance kinetics in a *Drosophila* model. Together, these results provide proof of concept for RNA phage surface engineering and establish a framework for the next generation of programmable viral nanoscaffolds.

## RESULTS AND DISCUSSION

### Identification of insertion-tolerant regions (ITRs) in the fiersphages

Insertion-tolerant regions (ITRs) are defined here as genomic positions that can accommodate sequence insertion without loss of phage viability. To identify such regions, we first introduced insertions into the genomic cDNA of MS2 and PP7 (10). As outlined schematically in Figure 1, MuA transposase was used in vitro to introduce a 1,131-bp MuA entranceposon randomly into plasmids carrying phage cDNA. After mutagenesis, each pool of insertion-containing DNA molecules was introduced into *E. coli* HB101 together with a compatible pJN105 plasmid expressing the cognate RNA replicase (RP) (19). Because RP is a nonstructural protein in both phages, complementation in trans should permit recovery of insertions that disrupt RP protein function, while still excluding insertions that interfere with essential RNA-level functions such as genome packaging. The primary screen was based on the inability of individual transposon clones to produce cDNA-derived phage progeny, as judged by the absence of a halo around each colony, which serves as a hallmark of active phage production. The majority of the clones (4,427 out of 4,555 for MS2 and 2,156 out of 2,228 for PP7) were defective in phage production, most likely because the transposon had inserted into essential phage cDNA regions or into sequences required for plasmid integrity. A total of 4,427 MS2 and 2,156 PP7 *E. coli* clones were pooled, and plasmid DNA was isolated. From this material, only plasmids carrying the transposon insertion within phage cDNA were recovered. These plasmids were then digested with NotI, self-ligated, and reintroduced into HB101. The resulting clones (3,575 for MS2 and 2,250 for PP7) were subjected to a second screen for their ability to produce functional phage particles. This procedure identified 29 and 26 non-redundant ITRs that tolerated a 15-bp insertion, as described in Methods and summarized in Table S1. Figure 2 shows the genomic distribution of the identified ITRs in the coding regions of MS2 and PP7, together with their corresponding phage productivity based on efficiency of plaque formation (EOP) on susceptible indicator host strains. Notably, most MS2 cDNA clones carrying a 15-bp insertion, even within RP, produced fewer progeny than the wild-type phage, whereas some PP7 cDNA clones carrying a 15-bp insertion produced phage at levels comparable to those of the wild type (Figure 2B), including MP15, RP289, and RP291 (Table S1). The mechanistic basis for this difference between MS2 and PP7, as well as the structural and functional flexibility associated with certain ITRs, including those located in structural proteins such as MP, remains to be investigated in future work.

**Figure 1.**
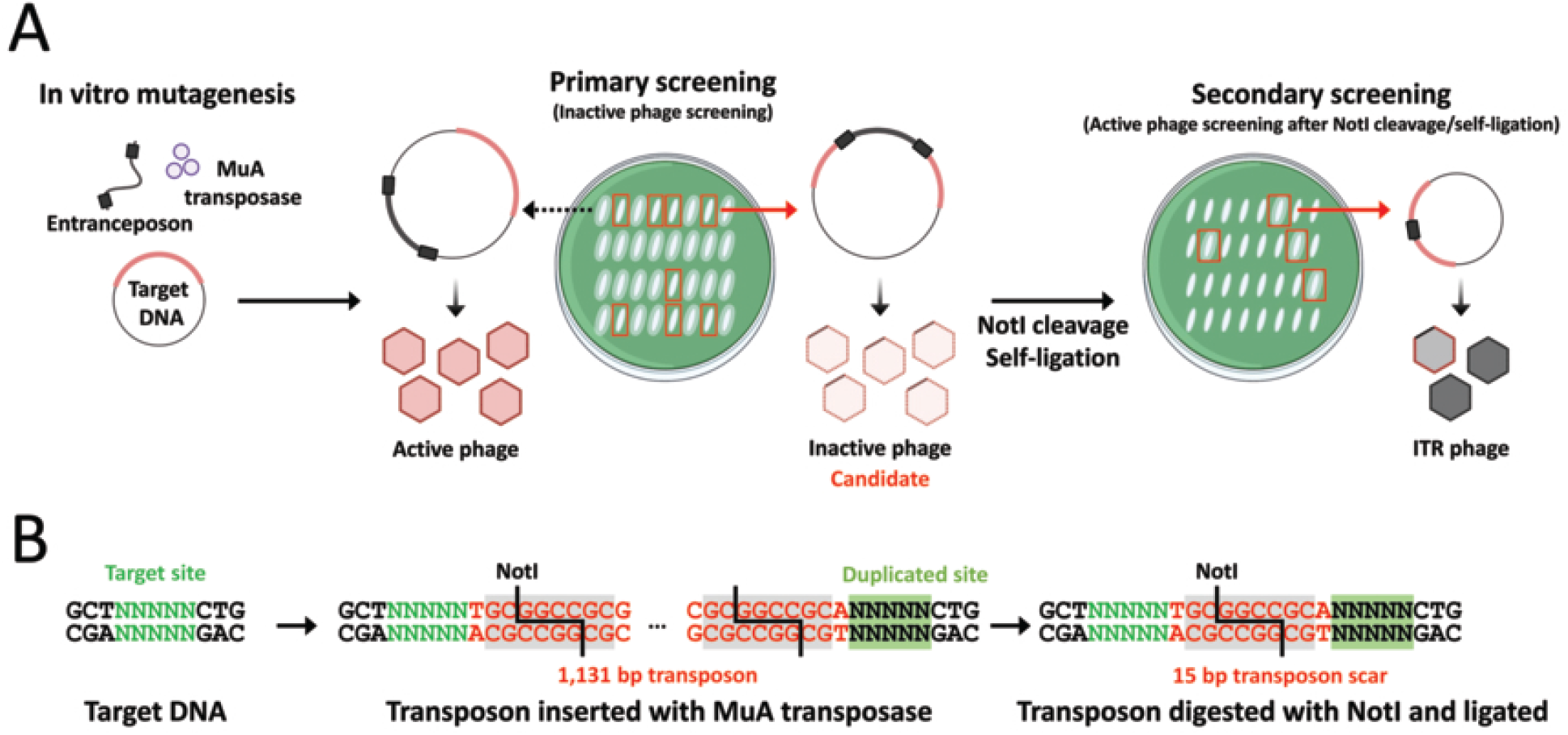
Strategy to Identify insertion-tolerant regions. **A**. Schematic overview of the in vitro transposon-based workflow used to identify insertion-tolerant regions (ITRs) in RNA phage genomes. Target DNA containing phage cDNA (colored) was subjected to in vitro mutagenesis with MuA transposase to generate random insertion of an entranceposon, resulting in insertion of a 1,131-bp DNA segment either within or outside the phage cDNA. Clones that initially failed to produce phage were collected, pooled, and subjected to plasmid isolation. The pooled plasmids were then digested with NotI and ligated to remove the transposon, leaving a 15-bp scar at each insertion site. The ligated plasmids were introduced into *E. coli*, and the resulting colonies were screened a second time for restoration of active phage production. **B**. Nucleotide sequences showing the transposon target site (TN_5_C), the flanking regions of the transposon insertion including the two NotI and duplicated target (N_5_) sites, and the 15-bp scar (TGCGGCCGCAN_5_) remaining after transposon removal.

**Figure 2.**
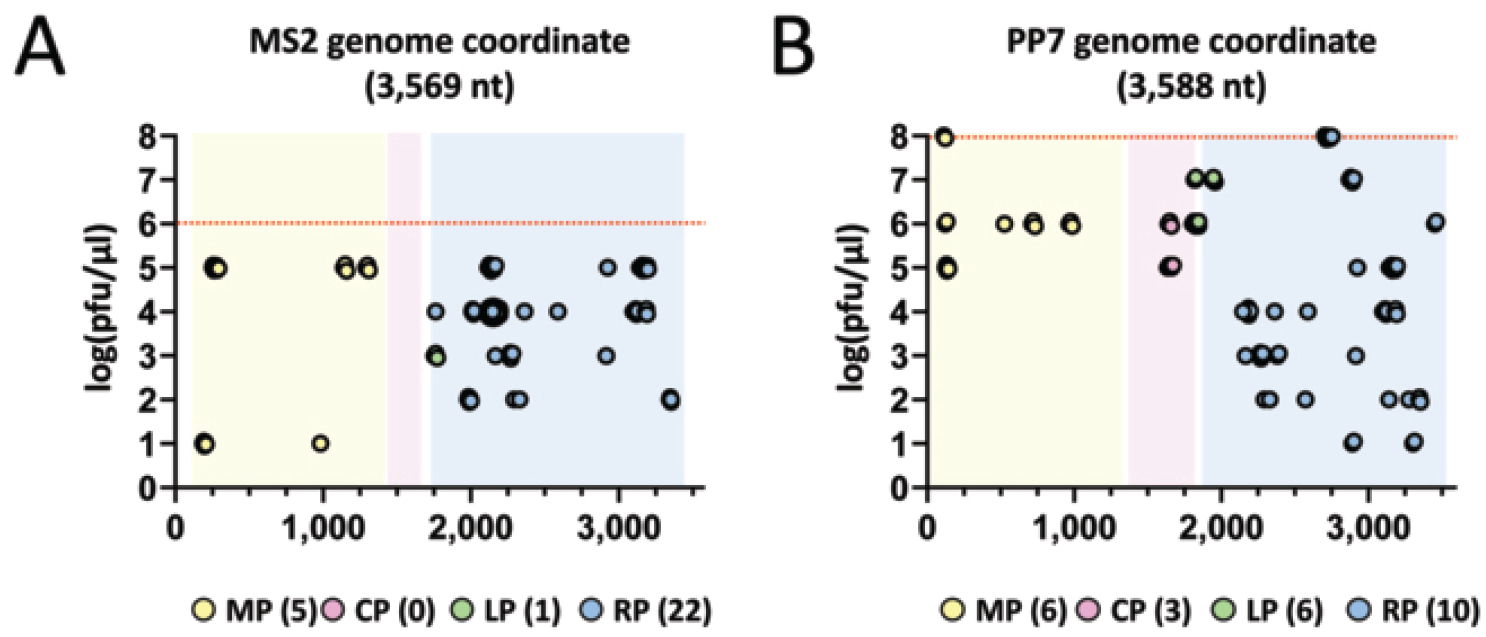
ITR profiles from PP7 and MS2. **A** and **B**. Genome-wide distribution of ITRs in the coding regions across the MS2 genome (3,569 nt) (**A**) and the PP7 genome (3,588 nt) (**B**). Infectious output is plotted as log(pfu/μl) based on culture supernatants from each surrogate strain expressing RP proteins. The coding regions for maturation protein (MP), coat protein (CP), and RNA-dependent RNA polymerase (RP) are color-shaded in the plot. Numbers in parentheses indicate the number of the non-redundant ITRs identified within each coding region. Red dotted vertical lines indicate the log(pfu/μl) values obtained from surrogate strains carrying the wild-type cDNA.

Most ITRs were located in the RNA replicase (RP) gene, particularly in MS2, in which 22 of 28 ITRs mapped to RP, with only 5 ITRs in MP and one in the lysis protein (LP). By contrast, the ITR distribution in PP7 was broader: among the 25 identified ITRs, 10 were located in RP, 6 in MP and LP, and 3 even within the coat protein (CP). It is especially notable that 3 ITRs were identified in the CP region of PP7, whereas no such permissive sites were detected in the CP of MS2. This observation is consistent with recent reports suggesting that PP7 and its capsid are more tolerant of modification than those of closely related RNA phages (20, 21). During our assessment of the pharmacokinetic (PK) properties of our in-house phages, we found that PP7 and MS2 exhibited superior in vivo stability in a *Drosophila* model, relative to other DNA phages, in which phages are administered through the fly medium (22) (Figure S1). These findings support the view that PP7 virions may provide a stable and compact scaffold for in vivo payload delivery applications.

### Characterization of the ITRs in CP

For further analysis, we selected the 3 ITRs identified in the PP7 CP region, given that CP is the major structural component of the capsid and is therefore the most relevant target for surface peptide display. A PP7 virion is composed of 90 CP protomers arranged as dimers together with one MP dimer (12, 23), making CP a more favorable scaffold than MP for robust peptide display. Structural modeling of the PP7 virion indicated that the ITRs at CP73 and CP74 are located in external surface loops or flexible, unstructured regions of the CP, whereas the ITR at CP82 is buried more deeply within the structural shell of the capsid (Figure 3A and B). Although CP73 and CP74 exhibited phage productivity approximately 2 logs lower than that of the wild-type phage, they nevertheless produced substantially more progeny than CP82 (Figure 3C).

**Figure 3.**
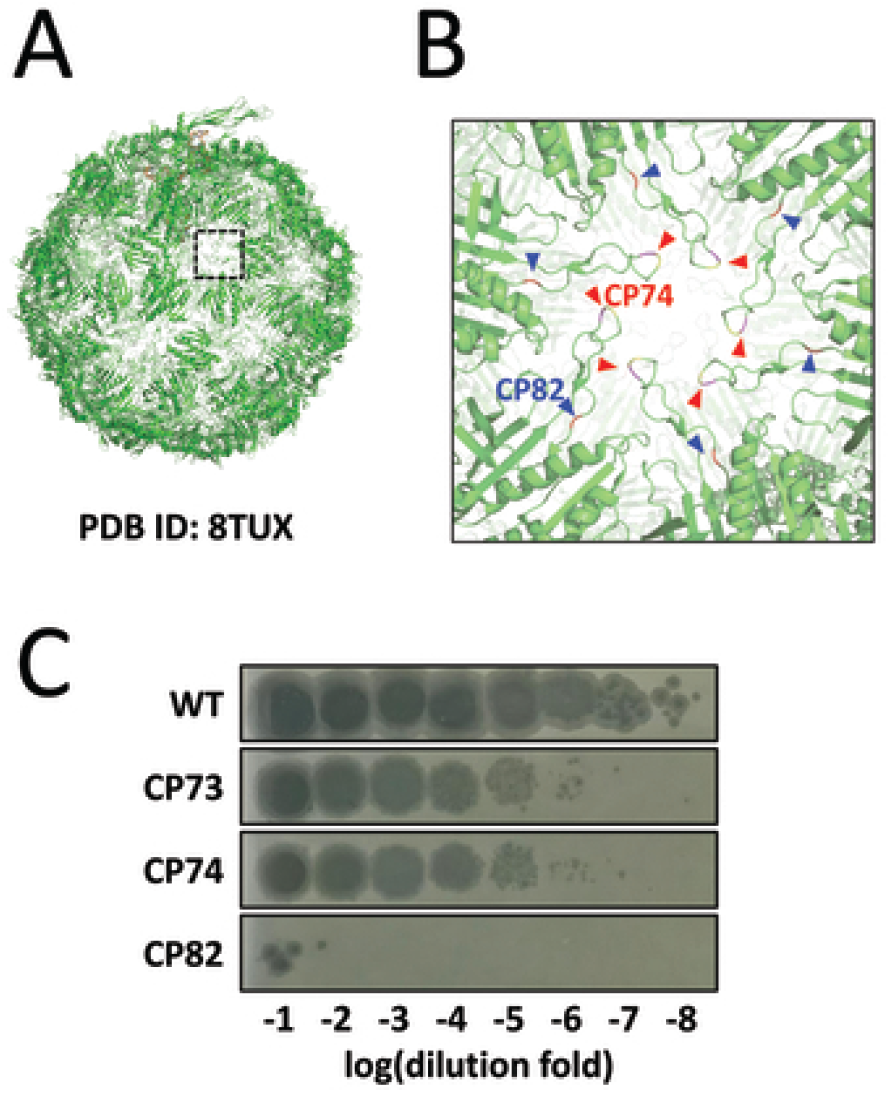
Characterization of ITRs in CP of PP7. **A**. Structural modeling showing the positions of ITRs identified within the PP7 CP. ITRs at CP73, CP74, and CP82 were mapped onto the PP7 capsid structure (PDB ID: 8TUX) to visualize their spatial positions. The boxed region corresponds to the capsomer area enlarged in panel **B**. **B**. Spatial localization of the ITRs within the capsomer. Red arrowheads indicate the 6 positions corresponding to CP73 and CP74 in flexible, surface-exposed loops, whereas blue arrowheads indicate the 6 positions corresponding to CP82 in deeper loop regions of the capsomer. **C**. Quantification of phage production from the CP ITR clones. Culture supernatants collected after 12 h of broth culture from surrogate strains carrying wild-type (WT) cDNA or one of the CP ITR clones (CP73, CP74, or CP82) were serially diluted 10-fold and assayed for plaque formation on PAO1ΔRF lawns. Numbers indicate log(dilution fold) values.

### Creation of the RNA phage particles tagged with a small peptide (RGD)

Because the CP73 and CP74 ITRs tolerated a 15-nt insertion, corresponding to 5 amino acids, we next attempted to introduce a small functional peptide tag into these positions. As both ITRs are located near the external loop region of the hexameric capsomer (Figure 3B), we inserted a 9-bp sequence encoding the Arg-Gly-Asp (RGD) motif at the CP74 ITR. RGD peptides are widely used ligands that mediate biological interactions with surface-exposed integrins in eukaryotic cells (24). In particular, the RGD motif can promote phagocytosis by binding RGD-recognizing integrins on professional phagocytes such as macrophages and dendritic cells. This interaction strengthens tethering and induces outside-in signaling, which in turn reorganizes actin to drive formation of the phagocytic cup and engulfment (25). To evaluate this concept in PP7, we constructed cDNA clones encoding RGD either at the CP74 position or at the MP149 position as a control (Figure 4A). Phage productivity was then assessed for cDNA clones expressing either RGD-tagged CP or RGD-tagged MP. The EOP of CP(RGD) was reduced by approximately 2 logs relative to that of the wild type, whereas the EOP of MP(RGD) was reduced by approximately 4 logs (Figure 4B). We further measured RNA genome copy numbers in the phage preparations as well as intracellular RNA levels in producer cells carrying the corresponding cDNA constructs. As shown in Figure 4C the genome copy number in CP(RGD) phage preparations was reduced by approximately 2 logs relative to that of the wild type, whereas intracellular RNA synthesis did not differ significantly from the wild-type level. These results suggest that RNA packaging is impaired in CP(RGD) phages. Because CP is directly involved in selective RNA packaging and virion assembly, it is likely that insertion of the RGD motif alters CP-RNA interactions in a manner that compromises RNA encapsidation without destabilizing the RNA genome itself. Accordingly, the reduced progeny yield of CP(RGD) is unlikely to result primarily from diminished infectivity.

**Figure 4.**
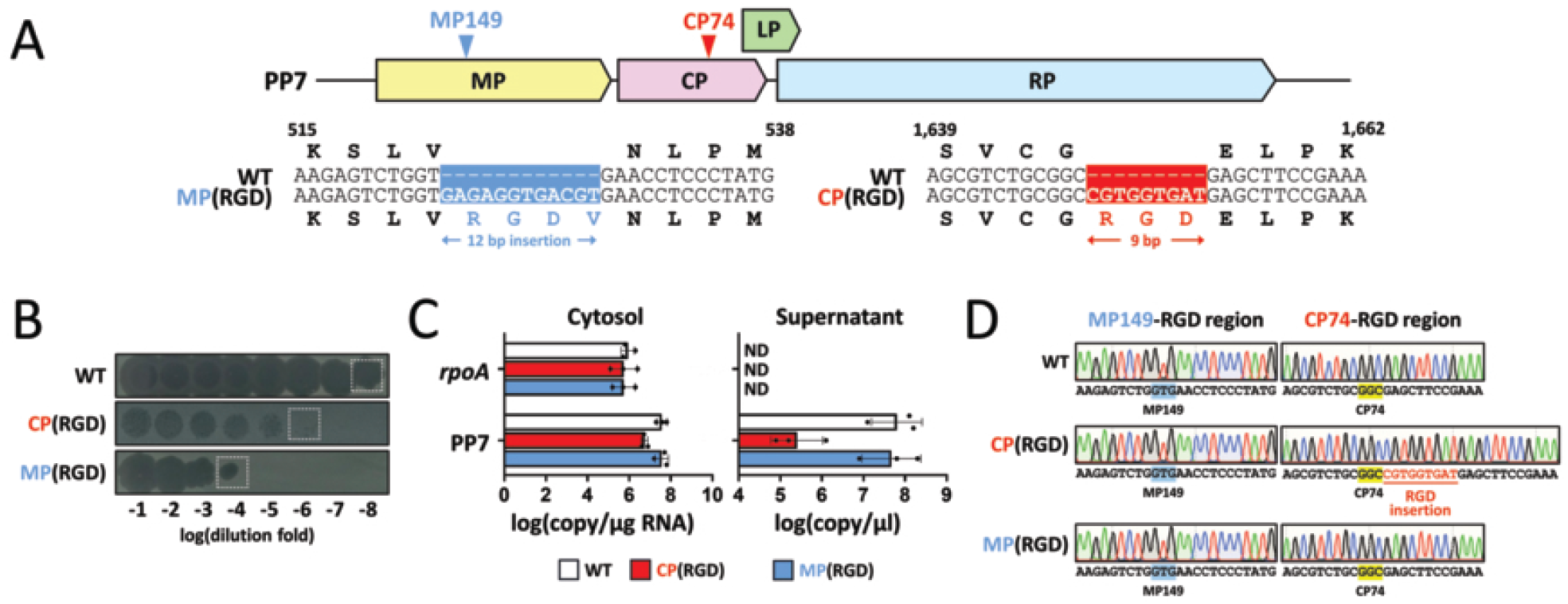
Engineering PP7 CP by tagging RGD peptide at C1650. **A**. Design of PP7 capsid mutants containing an RGD peptide insertion. Two insertion strategies were used: an RGD-encoding sequence was inserted either at the MP149 ITR [MP(RGD)] or at the CP74 ITR [CP(RGD)]. Corresponding nucleotide and amino acid sequences of the wild-type (WT) and engineered regions are shown, highlighting the inserted RGD motif and the insertion lengths: a 12-bp insertion to retaining the reading frame in MP(RGD) and a 9-bp insertion in CP(RGD). **B**. Quantification of phage production from PP7 capsid mutants containing RGD. Culture supernatants collected after 12 h from surrogate strains carrying WT cDNA or one of the RGD-tagged constructs [CP(RGD) or MP(RGD)] were 10-fold serially diluted and assayed for plaque formation on PAO1ΔRF lawns. Numbers indicate log(dilution fold) values. Dotted boxes indicate the plaque samples used for sequence analysis in panel **D**. **C**. RT-qPCR analysis of PP7 genomic RNA (gRNA) in cytosolic samples (left) and supernatant samples (right) from PP7 capsid mutants containing RGD. *rpoA* mRNA was used as a control (empty bars), as described in Methods. ND, not detected. Data represent three independent biological replicates with individual data points shown; error bars indicate standard deviation. **D**. Sequence verification of phage samples derived from PP7 capsid mutants containing RGD. Phage gRNA isolated from the plaques shown in panel **B** (dotted boxes) was analyzed by RT-PCR. Partial nucleotide sequences of the PCR products are shown in the electropherograms, including reversion mutations observed in MP(RGD).

It is also noteworthy that RNA levels in both phage particles and producer cells carrying MP(RGD) were comparable to those of the wild type, suggesting that MP(RGD) does not impair either genome synthesis or virion assembly. The lower EOP observed for MP(RGD) in Figure 4B is therefore most likely attributable to reduced infectivity, consistent with the direct role of MP in receptor interaction with the type IV pilus receptor (26). We have previously observed that critical point mutations can revert during phage propagation at a frequency of approximately 10^−4^ under our experimental conditions (27, 28),, which is consistent with the reported mutation rate of Qβ (∼1.4 × 10^−4^) (29, 30). This prompted us to ask whether the reduced EOP of MP(RGD) might reflect reversion events that specifically removed the insertion. To test this possibility, plaques indicated in Figure 4B were isolated and the nucleotide sequences of their RNA genomes were determined. Figure 4D shows representative sequence electropherograms spanning the RGD insertion sites. No sequence changes were detected in CP(RGD). By contrast, MP(RGD) phages carried reversion mutations that specifically eliminated the 12-bp insertion at the MP149 position, indicating that the MP149 ITR is sensitive to insertion sequence and/or insertion size, given that the 15-bp scar insertion caused only a 2-log reduction in phage production (Table S1). These findings further suggest that the CP74 ITR is dispensable for infectivity but contributes to efficient RNA packaging and/or virion assembly.

### Characterization of the RGD-tagged RNA phage particles

We next performed a more detailed characterization of CP(RGD) phage particles using large-scale phage preparations. First, we examined virion morphology by transmission electron microscopy (Figure 5A). Negative staining showed that CP(RGD) particles were morphologically indistinguishable from wild-type phages and retained packaged RNA genomes within the particles. We also analyzed CP identity in both phage preparations by mass spectrometry. As shown in Figures S2 and 5B and C, the mass spectral profiles confirmed the presence of the expected CP species in CP(RGD) phages. Together, these results demonstrate that CP(RGD) phages were successfully generated, although the reduced efficiency of RNA packaging or virion assembly resulted in an approximately 100-fold lower particle yield.

**Figure 5.**
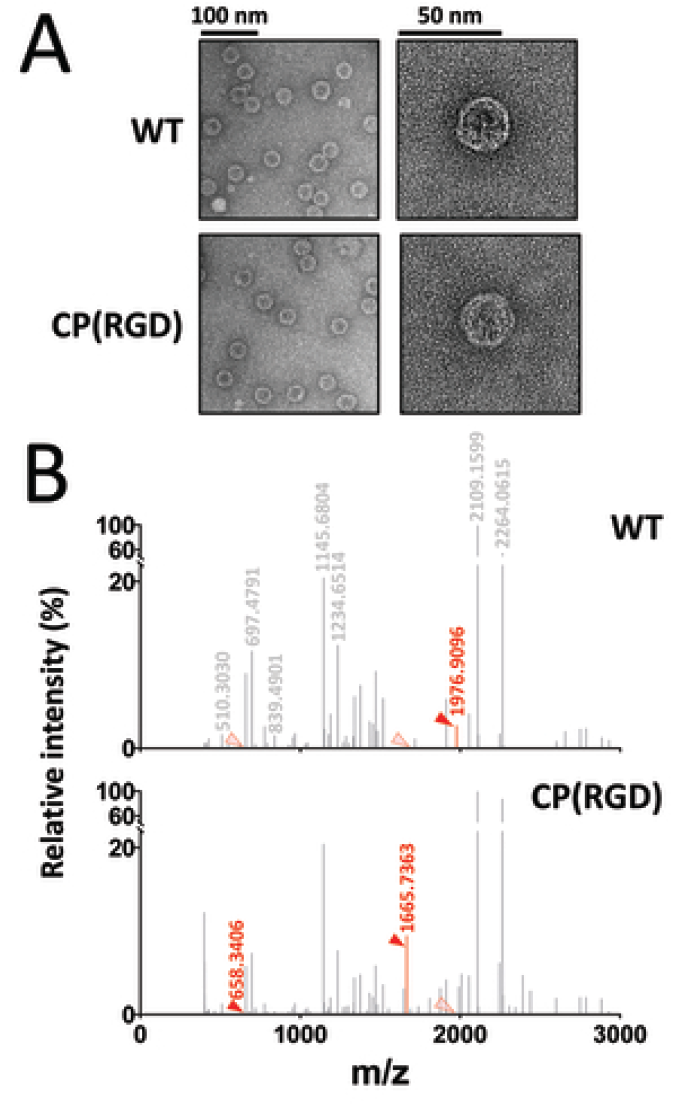
Validation of engineered phage particles. **A**. Representative transmission electron micrographs of wild-type (WT) and CP(RGD) phage particles. Images are shown at two magnifications with scale bars of 100 nm and 50 nm to compare particle morphology and size. **B**. Mass-spectrometric validation of CP tagging. Relative signal intensities of detected peptide ions are shown for WT and CP(RGD) phage particles. Peaks corresponding to peptide fragments that differ between WT and CP(RGD) are indicated by arrowheads (see Figure S2), providing molecular confirmation of capsid engineering.

Prior to assessing the functional effect of RGD display on in vivo pharmacokinetic (PK) behavior, we first examined the in vitro stability of CP(RGD) phage particles in phage buffer at room temperature. Wild-type and CP(RGD) phages showed no detectable difference in stability for up to 48 h (Figure S3). We then evaluated the in vivo stability of CP(RGD) in comparison with that of the wild type using a *Drosophila* PK model. As shown in Figure 6, the in vivo persistence of CP(RGD) was substantially reduced relative to that of the wild type, with complete clearance by 36 h after administration, whereas the wild-type phage remained detectable up to 48 h. These results suggest that surface display of RGD on the major capsid protein enhances phagocytic clearance in *Drosophila*. It will be important in future studies to determine whether RGD display also affects feeding efficiency and/or gastrointestinal uptake in this model. More importantly, validation in larger animal models such as mice or rats will be required to define the functional effects of RGD tagging on phage pharmacokinetics in greater detail.

**Figure 6.**
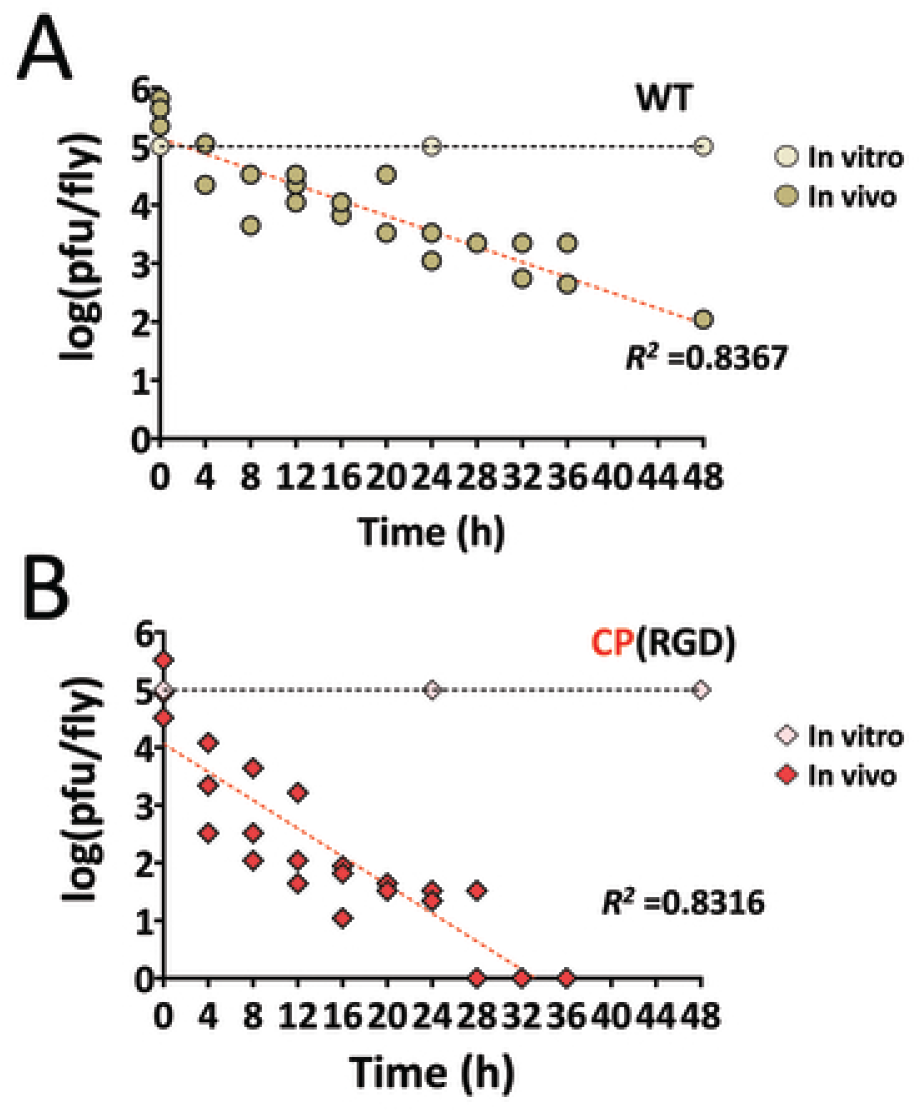
Stability of engineered phage particles in *Drosophila*. In vivo stability of WT (**A**) and CP(RGD) (**B**) phage particles. Phage titers, expressed as log(pfu/fly) were determined over time from fly homogenates following the scheme shown in Figure S1A. Slopes represent the in vivo stability of the administered phages, and the corresponding *R*^*2*^ values are indicated. For comparison, in vitro stability data for both phages (Figure S3) are shown as the vertical lines relative to the initial value set to 10^5^ for comparison. Data are from three independent biological replicates, with individual data points displayed.

## Conclusions

In this study, we established a genome-wide workflow for identifying insertion-tolerant regions (ITRs) in two fiersphages, MS2 and PP7, using MuA transposase-mediated in vitro insertion followed by functional screening for progeny production. This approach enabled recovery of nonredundant sets of ITRs that tolerate a 15-bp insertion, yielding 28 ITRs in MS2 and 25 ITRs in PP7. Although most insertions reduced phage productivity, the tolerance landscapes of the two phages differed substantially: MS2 ITRs were concentrated predominantly in RP, whereas PP7 exhibited a broader distribution across RP, MP, LP, and notably CP. These results demonstrate that insertion tolerance is highly phage specific and cannot be predicted solely from relatedness, emphasizing the importance of direct experimental ITR mapping.

A key outcome of this work is the identification and functional analysis of CP ITRs in PP7, which directly expands the practical engineering space for phage surface display. Structural mapping indicated that CP73 and CP74 are located in external and/or flexible regions of the capsid, whereas CP82 lies deeper within the shell, consistent with the observed differences in phage productivity. Leveraging CP74 as a display-compatible ITR, we inserted an RGD motif as a proof-of-concept ligand. The resulting CP(RGD) particles retained native morphology and the expected coat-protein identity, and RNA synthesis in producer cells remained intact, whereas RNA packaging and/or assembly efficiency was reduced, as reflected by lower particle yield and lower genome copy number in purified virions. By contrast, insertion of the same motif into MP reverted rapidly through excision of the insert, indicating that certain ITRs may be sensitive to insertion sequence and insertion size in ways that affect essential stages of the viral lifecycle. Collectively, these findings distinguish ITRs that preserve infectivity from those that primarily modulate packaging and assembly efficiency, and they demonstrate that insertion tolerance should be evaluated across multiple functional dimensions rather than on viability alone.

Looking forward, several directions emerge that could strengthen both the mechanistic understanding and the translational utility of ITR-guided fiersphage engineering. First, the striking divergence between MS2 and PP7, particularly the presence of CP ITRs only in PP7, warrants targeted investigation into the structural and RNA-level determinants that shape genome flexibility, including local RNA structures, packaging signal density, and CP-RNA interactions. Second, because RP was supplied in trans in the present study, ITRs in the RP region should be revisited to disentangle RNA-level constraints from protein-level requirements. Third, engineering of CP74 showed that surface display can substantially alter in vivo behavior: CP(RGD) showed accelerated clearance in *Drosophila*, consistent with enhanced phagocytic recognition, while remaining stable in vitro. This finding motivates systematic studies of ligand identity, insertion length, linker architecture, and display density to separate effects on packaging and assembly from effects on PK properties. It also underscores the need for expanded validation in mammalian models to define biodistribution, clearance pathways, and immune interactions under more clinically relevant conditions. Finally, comprehensive ITR maps could serve as standardized “landing-pad” catalogs for modular genome manipulation, enabling barcoding, multisite insertion, and rational virion functionalization, provided that insertion tolerance is defined quantitatively in terms of yield, packaging efficiency,infectivity, and genetic stability, and is accompanied by design rules that generalize across phages. Together, these results suggest that PP7 may serve as a more predictable and engineerable RNA nanoplatform for both mechanistic RNA virology and delivery-oriented applications.

## ACKNOWLEDGEMENTS

This work was supported by the National Research Foundation of Korea (NRF) Grants (RS-2022-NR070540 and RS-2022-NR067344).

## AUTHOR CONTRIBUTIONS

Y.-H.C. conceived and designed the research. S.-Y.C., M.-J.K., and O.H.K. designed and performed the experiments and collected and analyzed the data. C. P., H.-W.B., and H.C. provided reagents and critical comments. S.-Y.C., M.-J.K, H.-W.B, and Y.-H.C. wrote the manuscript. All authors reviewed and approved the final version of the manuscript.

## METHODS

### Bacterial strains and culture conditions

The bacterial stains used in this study are listed in Table S2. *Pseudomonas aeruginosa* strains (PAO1 and PAK and their derivatives lacking R and F pyocins) and *Escherichia coli* strains (DH5α, HB101, C3000, and SM10(λ*pir*)) were routinely cultured in Luria-Bertani (LB) broth (1% tryptone, 0.5% yeast extract, 1% NaCl) or on LB agar plates containing 2% bacteriological agar (MBcell). Cetrimide agar (Difco) was used for selection of *P. aeruginosa* strains. All strains harboring phage genes were incubated at 30°C. Where appropriate, antibiotics were added at the following concentrations (μg/ml): gentamicin (50), carbenicillin (200), and kanamycin (200) for *P. aeruginosa*; gentamicin (25), ampicillin (50), and kanamycin (50) for *E. coli*.

### Preparation of phage lysates

Phages were prepared from culture supernatants of surrogate strains harboring phage cDNA (HB101 for MS2 and PAK for PP7). Cells were grown in LB broth for 12-18 h at 30°C. NaCl (1 M) and chloroform (0.1% v/v) were then added to the culture supernatants, and phage lysates were obtained by centrifugation. Phage particles were precipitated with 10% polyethylene glycol 8,000 (Sigma-Aldrich), pelleted by centrifugation, and resuspended in phage buffer (0.1 M NaCl, 10 mM MgSO_4_·7H_2_O, 50 mM Tris-Cl, pH 7.5). Phage samples were treated with DNase I (100 μg/ml; Sigma-Aldrich) and RNase A (50 μg/ml; Thermo Fisher Scientific) for 1 h and then extracted with chloroform. Phage particles were ultracentrifuged at 300,000 × *g* for 4 h and resuspended in phage buffer to achieve titers exceeding 10^11^ pfu/ml.

### In vitro mutagenesis and ITR screen

In vitro mutagenesis of miniTn*7*-based MS2 and PP7 cDNA constructs used as target DNA (Table S2) was performed using MuA transposase (Thermo Fisher Scientific). The transposition reaction was carried out with 320 ng of target DNA for 1 h at 30°C and terminated by incubation at 75°C for 10 min. Appropriately diluted reaction mixtures were introduced into HB101 by electroporation, which expresses pJN105-borne RNA replicase (either pJN-RP_MS2_ or pJN-RP_PP7_) (Table S2). The resulting colonies (4,555 for MS2 and 2,228 for PP7) were first screened for their inability to produce phage on lawns of the corresponding indicator strains (i.e., C3000 for MS2 and PAO1ΔRF for PP7). Candidate clones (4,427 for MS2 and 2,156 for PP7) were pooled and subjected to plasmid isolation. The recovered plasmids were digested with NotI, ligated with T4 DNA ligase, and reintroduced into HB101 containing either pJN-RP_MS2_ or pJN-RP_PP7_ by electroporation. The resulting colonies (3,575 for MS2 and 2,250 for PP7) were then subjected to a second screen for restoration of active phage production on the appropriate indicator lawns. The candidate clones (155 for MS2 and 98 for PP7) were further confirmed to produce phage progeny and final candidate clones (128 for MS2 and 72 for PP7) were obtained. These cDNA clones were expected to harbor phage cDNA containing 15-bp random insertions at ITRs. PCR using two primer pairs including NotI-mini (Table S3) was carried out to determine the approximate positions of the ITRs.

### Quantification of phage production

Phage production was quantified by plaque assay for pfu measurement and by RT-qPCR following reverse transcription for genome copy number assessment. For plaque assays, appropriately diluted phage samples (10 μl) were mixed with 3 ml of 0.7% top agar containing ∼10^8^ indicator cells (PAO1ΔRF for PP7 and C3000 for MS2) and overlaid onto LB agar plates. Plaque formation was examined after incubation at 30°C for 16-24 h (31). Phage genome copy numbers were determined as previously described (10) using phage-containing supernatants from cDNA-carrying PAKΔRF cells. Phage RNA was isolated from phage samples using TRIzol (Invitrogen) according to the manufacturer’s instructions. To measure the cytosolic RNA levels, cells were grown to mid-exponential growth phase at 30°C, and total RNA was extracted using the RNeasy Mini Kit (Qiagen). All RNA samples were treated with DNase I (Qiagen) and subjected to RT-qPCR with primers listed in Table S3 using the StepOnePlus Real-Time PCR System (Applied Biosystems). Phage genome copy numbers were determined from standard curves in comparison with the *rpoA* mRNA levels, as previously described (32).

### RGD tagging and validation

RGD tagging was introduced by 4-primer splicing by overlap extension PCR followed by sequence- and ligation-independent cloning into a HindIII-digested miniTn*7* vector, as described previously (33, 34) (Table S2). The resulting constructs were introduced into the surrogate strain PAKΔRF by conjugation to generate the engineered phage particles carrying the RGD insertion. These engineered phage particles were validated by transmission electron microscopy (TEM) and mass spectrometry. For TEM,formvar-coated TEM grids were surface-treated for 10 min and then floated on 10 μl of 1:100 diluted phage samples. Negative staining was performed by applying 5 μl of 2% (w/v) uranyl acetate solution (pH 4.0) for 15 s. Excess stain was removed, and the grids were air-dried for 5 min. Samples were imaged using a 120 kV Tecnai transmission electron microscope at magnifications ranging from ×500,000 to ×120,000. For mass spectrometry, phage proteins were resolved by 15% SDS-PAGE and stained with Coomassie Brilliant Blue. CP bands were excised and subjected to in-gel tryptic digestion (20 ng/μl). The digested peptides were dried under vacuum and analyzed by MALDI-TOF mass spectrometry performed in positive ion mode over a mass range of 500-300,000 Da using a nitrogen laser (337 nm) with delayed extraction, as described elsewhere (35).

### *Drosophila* experiments

In vivo stability of phage particles was assessed using a *Drosophila* feeding model (22). Briefly, *Drosophila melanogaster* strain Oregon R was grown and maintained at 25°C using the corn meal-dextrose medium containing 6.24% dry yeast, 4.08% corn meal, 8.62% dextrose, 0.93% agar, 10% tegosept, and 0.45% (v/v) propionic acid. To evaluate phage pharmacokinetics in *Drosophila*, male flies were fed for 16 h on corn meal-dextrose medium overlaid with 10^10^ pfu of phage particles and then transferred to phage-free medium. At 4-h intervals, groups of three flies were collected and homogenized in 100 μl of phage buffer. The homogenates were used for pfu measurement.

### Structural modeling of PP7 capsid

Structural modeling was performed using AlphaFold3 to generate three models for each construct, and the highest-ranked model (ranked_0.pdb), selected on the basis of the average predicted local distance difference test (pLDDT) score, was used for subsequent analysis. Predicted models were examined in a hexameric configuration, visualized with PyMOL, and compared with the experimentally determined PP7 capsid structure (PDB ID: 8TUX).

### Statistics

Data are presented as means together with individual data points and were analyzed using GraphPad Prism version 8.4.3.

### DATA AVAILABILITY

The data reported in this study will be made available by the corresponding author upon request. Additional information required to reanalyze the reported data is also available upon request.

